# *Clostridioides difficile* binary toxin binding component (CDTb) increases virulence in a hamster model

**DOI:** 10.1101/2021.08.03.454966

**Authors:** Morgan Simpson, Terry Bilverstone, Jhansi Leslie, Alexandra Donlan, M Jashim Uddin, William Petri, Natasha Marin, Carsten Schwan, Sarah Kuehne, Nigel P. Minton, William A. Petri

## Abstract

*Clostridioides difficile* is the leading cause of hospital-acquired gastrointestinal infection, in part due to the existence of binary toxin (CDT)-expressing hypervirulent strains. We have previously shown that CDT interacts with the TLR2/6 heterodimer to induce inflammation, and in this study we further explore this interaction as well as the contribution of the separate components of CDT, CDTa and CDTb. We found that the binding component, CDTb, is capable of inducing inflammation. Additionally, complementation of a CDT-deficient *C. difficile* strain with CDTb alone restored virulence in a hamster model of *C. difficile* infection. Overall, this study demonstrates that the binding component of *C. difficile* binary toxin contributes to virulence during infection.

## Introduction

*Clostridioides difficile*, a gram-positive, spore-forming anaerobe, is the causative agent of *Clostridioides difficile* infection (CDI), a gastrointestinal infection typically characterized by high levels of inflammation and diarrhea. This bacterium is considered an urgent health threat by the CDC (1), and was shown in a 2015 study to be responsible for approximately 500,000 infections and 29,000 deaths (2). *C. difficile* typically infects those with dysbiosis, a state of disruption in the healthy intestinal microbiota leading to reduced and/or skewed microbial diversity, commonly induced through use of broad-spectrum antibiotics (3). This reduced microbial population and diversity allows *C. difficile* to establish a niche and begin toxin production. These toxins lead to disruption of the host intestinal epithelial barrier, production of pro-inflammatory cytokines, and recruitment of inflammatory immune cells to the site of infection (4). The host immune response to CDI is critical in determining patient outcome, as immune biomarkers have been shown to be more predictive of time to disease resolution than bacterial burden (5). While effective antibiotic treatment is available, 1 in 5 patients will experience recurrent infection (2), highlighting the need for further understanding of the host response to aid in the development of improved or novel therapeutics.

The past few decades have seen an overall increase in both the frequency and severity of CDI, a phenomenon that has been primarily associated with the emergence of hypervirulent strains of *C. difficile* (6). In addition to the primary virulence factors Toxin A and Toxin B, these strains express a third toxin called *C. difficile* transferase, or CDT (7). CDT is a binary toxin with ADP-ribosyltransferase activity and consists of an enzymatic component, CDTa, and a binding component, CDTb. Following the binding of CDTb to its host cell receptor, lipolysis-stimulated Lipoprotein receptor (LSR) (8), CDTa binds to CDTb and induces endocytosis into the host cell. Upon acidification of the endosome, CDTb inserts itself into the endosomal membrane and forms a pore through which CDTa escapes into the cytosol. CDTa then ADP-ribosylates actin, thereby preventing its elongation and leading to cytoskeletal disruption (7). CDT also induces the formation of microtubule protrusions at the apical surface which are thought to aid in bacterial adherence to host epithelial cells (9). In addition, previous work from our group demonstrated that CDT can suppress protective eosinophilic responses (10) and signal through TLR2 and TLR6 to induce downstream inflammation (11). Yet while the CDT holotoxin has been shown to contribute to the severity of CDI through the aforementioned responses, the exact contribution of its individual components, CDTa and CDTb, remain unknown.

In this study, we sought to investigate the role of the individual components of CDT in TLR2/6-mediated inflammation, as well as during *in vivo* infection. We found that the binding component CDTb is capable of inducing NF-κB activation via the TLR2/6 heterodimer and can induce IL-1β expression in bone marrow-derived dendritic cells (BMDCs) in a dose-dependent manner. In addition, we developed novel mutant strains of *C. difficile* capable of producing CDTa or CDTb alone to determine the contribution of CDTb during CDI. While expression of CDTb without CDTa could not induce significant disease in a mouse model of CDI, we observed severe disease and mortality in a hamster model. Overall, we have found that CDTb is capable of inducing downstream inflammation and enhancing virulence in a hamster model of infection.

## Materials and Methods

### Cell Culture and Toxins

HEKBlue-hTLR2 reporter cells were obtained from Invivogen and grown in Dulbecco’s Modified Eagle’s Medium (DMEM) supplemented with 4.5 g/L glucose, 2 mM L-glutamine, 10% fetal bovine serum, 100 U/mL penicillin, 100 μg/mL streptomycin, 100 μg/mL Normocin (Invivogen), and 1x HEK-Blue Selection (Invivogen). Cells were rinsed with warm phosphate-buffered saline, detached with a cell scraper, and resuspended at a density of 2.8 × 10^5^ cells/mL in HEK-Blue Detection media (Invivogen). Neutralizing antibodies against TLR2, TLR1, or TLR6 (Invivogen) were added to the cells at a concentration of 5 μg/mL and incubated for 1 hour at 37°C. Following the incubation, CDTa and/or CDTb (5 ng/mL each) was added. An equivalent volume of endotoxin-free water (20 μL;Fisher Scientific) was used as a negative control. Cells were incubated at 37°C in a 5% CO_2_ incubator for 6-16 hours. Secreted embryonic alkaline phosphatase (SEAP) was quantified by taking the optical density at 655 nm in a spectrophotometer.

Murine BMDCs were generated as previously described with minor modifications (10). Femurs and tibia were removed and spun within a microcentrifuge tube briefly at high speed to remove bone marrow. Cells were cultured in RPMI 1640 media (Gibco) containing 10% fetal bovine serum (FBS), 2 mM l-glutamine, and 100 U/mL penicillin and 100 U/mL streptomycin. Media were supplemented with 10 ng/mL GM-CSF (Peprotech) and cells isolated from one femur and tibia were seeded into a T75 vent cap tissue culture flask. Cells were cultured for 7 days and supplemented with fresh media on day 3. On day 7, BMDCs were detached from the flask and resuspended at 5 × 10^5^ cells/mL in fresh media. Cell suspension (540 μL) was added to each well of a 48-well tissue culture plate. Cells were stimulated with CDTa and/or CDTb as indicated at a volume of 60 uL. After an overnight incubation at 37 °C with 5% CO_2_, the supernatant was removed. Cells were treated with 600 μL Trizol (Ambion) for 1 min, then removed and frozen at −80 °C for later analysis.

Purified CDTa and CDTb were expressed in *Bacillus megaterium* and purified as described previously (9).

### Cytokine Detection

IL-1β and IL-6 were detected in tissue lysate using the Mouse IL-1β and IL-6 DuoSet ELISA kits (R&D Systems) according to the manufacturer’s instructions. Cytokine levels were normalized to cecal tissue weight. IL-1β transcript production from BMDCs was assessed using quantitative reverse transcription PCR. RNA was isolated via chloroform-phenol phase separation followed by processing via the RNeasy isolation kit (Qiagen). Contaminating genomic DNA was digested using the Turbo DNA-free kit (Ambion) and samples were run through the RNeasy kit once more to remove any remaining genomic DNA. RNA was reverse transcribed with the Tetro cDNA synthesis kit (Bioline) according to manufacturer’s instructions, and the resulting cDNA was purified using the Qiagen PCR purification kit. IL-1β gene expression was quantified by PrimePCR FAM Probe assay (Bio-rad) using SsoAdvanced Universal Probes Supermix (Bio-rad) using the Bio-rad two-step amplification protocol. Gene expression was normalized to the housekeeping gene Hprt. Hprt gene expression was quantified by PrimePCR SYBR Green primer assay (Bio-rad) using SsoAdvanced Universal SYBR Green Supermix.

### Generation of CDT Mutant Strains

Strains and plasmids used in this study are listed in Table 1, whilst primers are listed in Table 2. The genes encoding *cdtA* and *cdtB* were deleted using allelic-exchange (AE) technology (12). To achieve this, left and right homology arms, corresponding to the regions annealing immediately upstream and downstream of *cdtA/B*, were amplified by PCR using *cdtAB* LAF/RAR and *cdtAB* RAF/RAR primer sets respectively. The homology arms were then spliced together by splicing by overlap-extension (SOEing) PCR by means of their overlapping 20bp homologous regions before cloning the ensuing product into pMTL-YN4 using flanking *Sbfl-Ascl* restriction sites, thus generating the knockout cassette (KOC) pMTL-YN4-*cdtAB* KOC. The plasmid was then conjugated into *C. difficile R20291*Δ*pyrE* exactly as described previously and transconjugants were selected on the basis of thiamphenicol resistance (13). Thereafter, single cross-over integrants (SCOs) were identified by two parallel PCR screens using *cdtAB* diag F/ YN4 primers for left arm recombinants and YN4 F/ *cdtAB* diag R primers for right arm recombinants respectively (data not shown). To select for double cross-over recombinants, SCO integrants were harvested, diluted 1 ×l0^−3^ and cultured onto *Clostridium difficile* minimal medium (CDMM) (14) containing 500μg/ml 5-fluoroorotic acid (FOA) and 1μg/ml uracil, to force plasmid loss through the counter-selection marker *pyrE*, and to select for double cross-over mutants before confirming plasmid loss on the basis of thiamphenicol sensitivity. Deletion mutants were confirmed to be as intended by PCR analysis using *cdtAB* diag F/R primers. Indeed, the deletion mutant generated a circa 4kbp product compared with the 4.6kbp product of its wild-type counterpart (Figure 2A). Finally, the *pyrE* allele was restored to wild-type using pMTl-YN2 exactly as described previously (13).

**Table 1:**
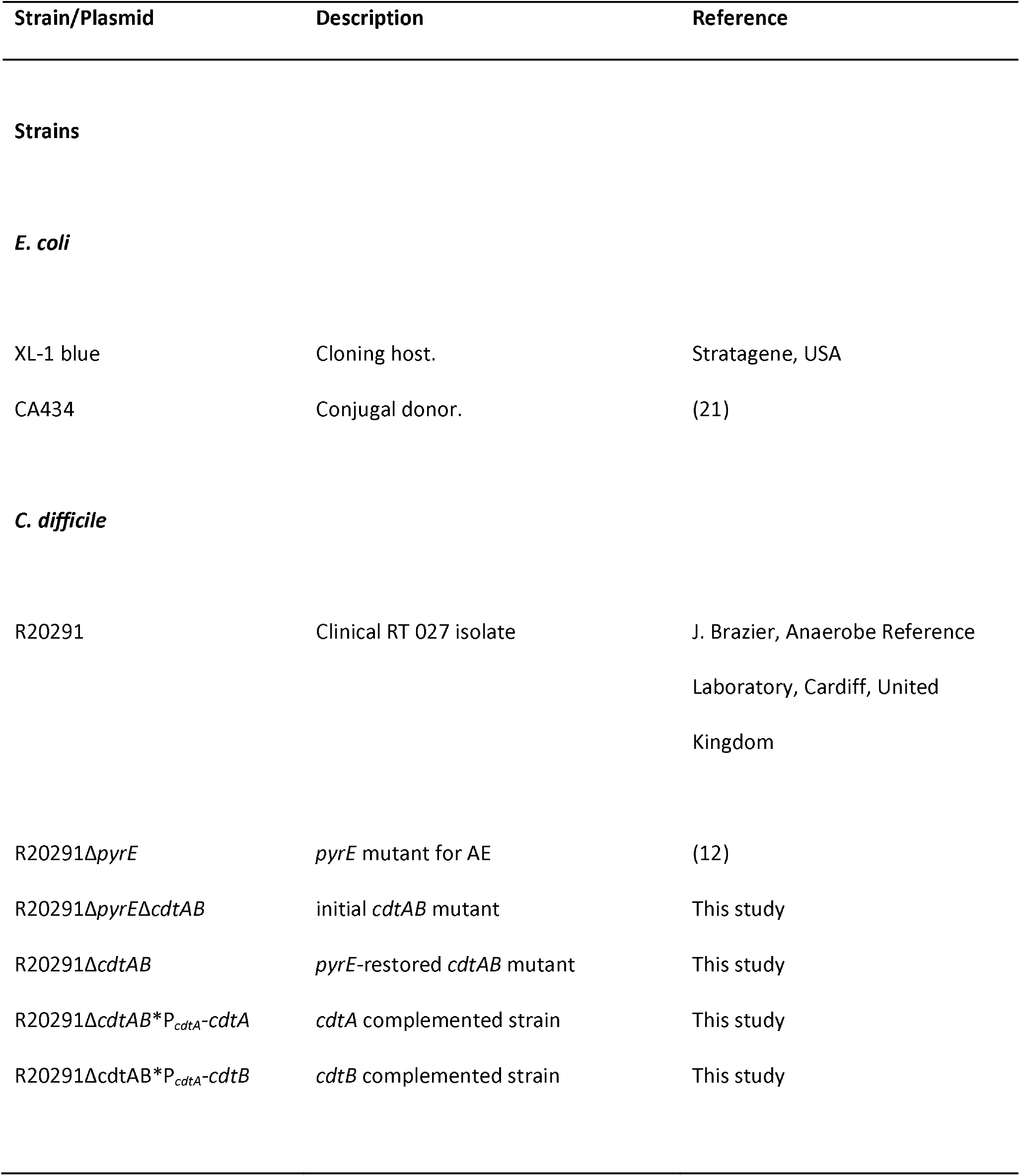
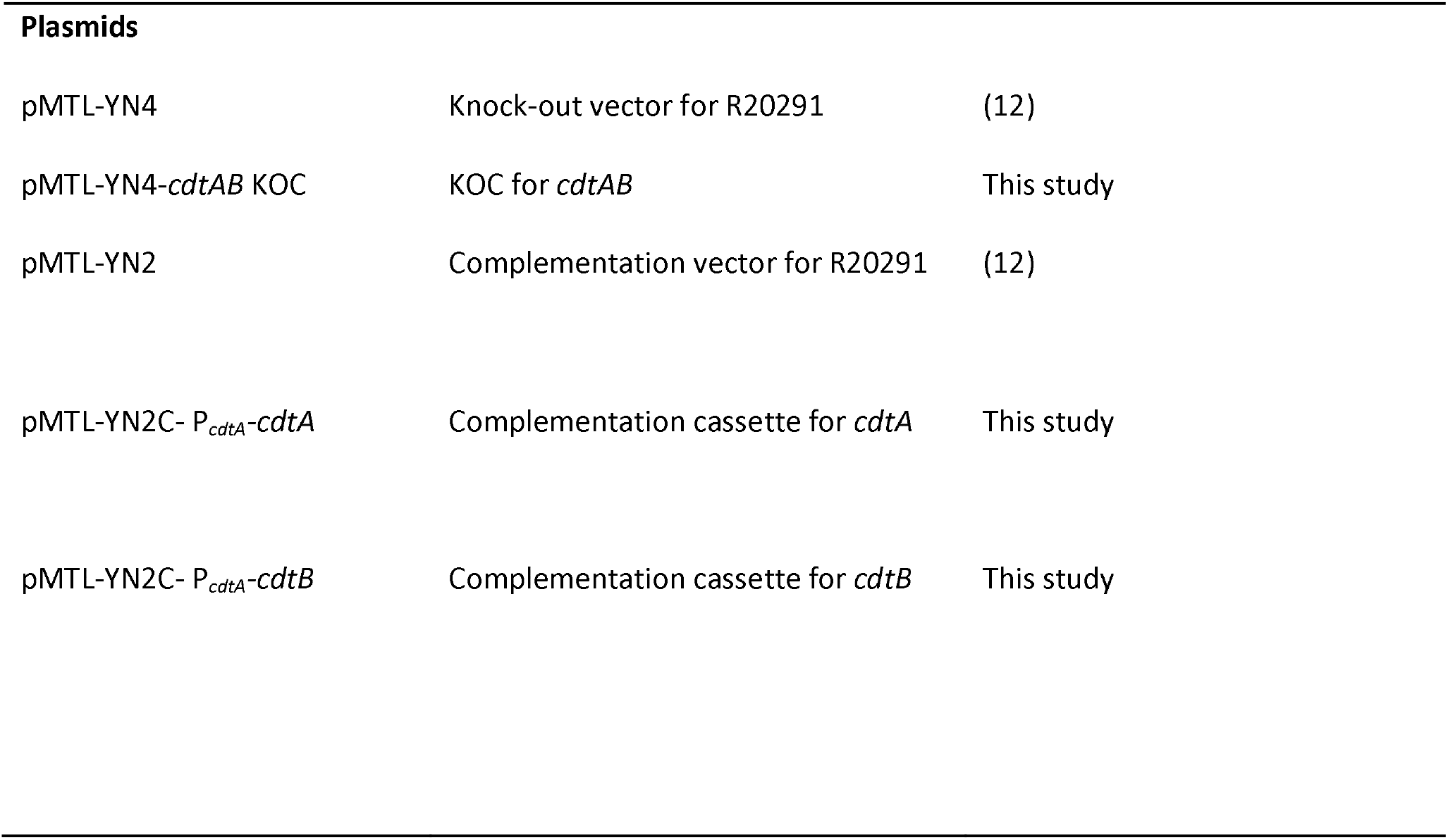
Strain and plasmids used in this study.

| Strain/Plasmid | Description | Reference |
| --- | --- | --- |
| **Strains** |  |  |
| *E. coli* |  |  |
| XL-1 blue | Cloning host. | Stratagene, USA |
| CA434 | Conjugal donor. | (21) |
| *C. difficile* |  |  |
| R20291 | Clinical RT 027 isolate | J. Brazier, Anaerobe Reference<br>Laboratory, Cardiff, United<br>Kingdom |
| R20291ΔpyrE | pyrE mutant for AE | (12) |
| R20291ΔpyrEΔcdtAB | initial cdtAB mutant | This study |
| R20291ΔcdtAB | pyrE-restored cdtAB mutant | This study |
| R20291ΔcdtAB*P<sub>cdtA</sub>-cdtA | cdtA complemented strain | This study |
| R20291ΔcdtAB*P<sub>cdtA</sub>-cdtB | cdtB complemented strain | This study |
| **Plasmids** |  |  |
| pMTL-YN4 | Knock-out vector for R20291 | (12) |
| pMTL-YN4-*cdtAB* KOC | KOC for *cdtAB* | This study |
| pMTL-YN2 | Complementation vector for R20291 | (12) |
| pMTL-YN2C- P<sub>*cdtA*</sub>-*cdtA* | Complementation cassette for *cdtA* | This study |
| pMTL-YN2C- P<sub>*cdtA*</sub>-*cdtB* | Complementation cassette for *cdtB* | This study |

**Table 2:**
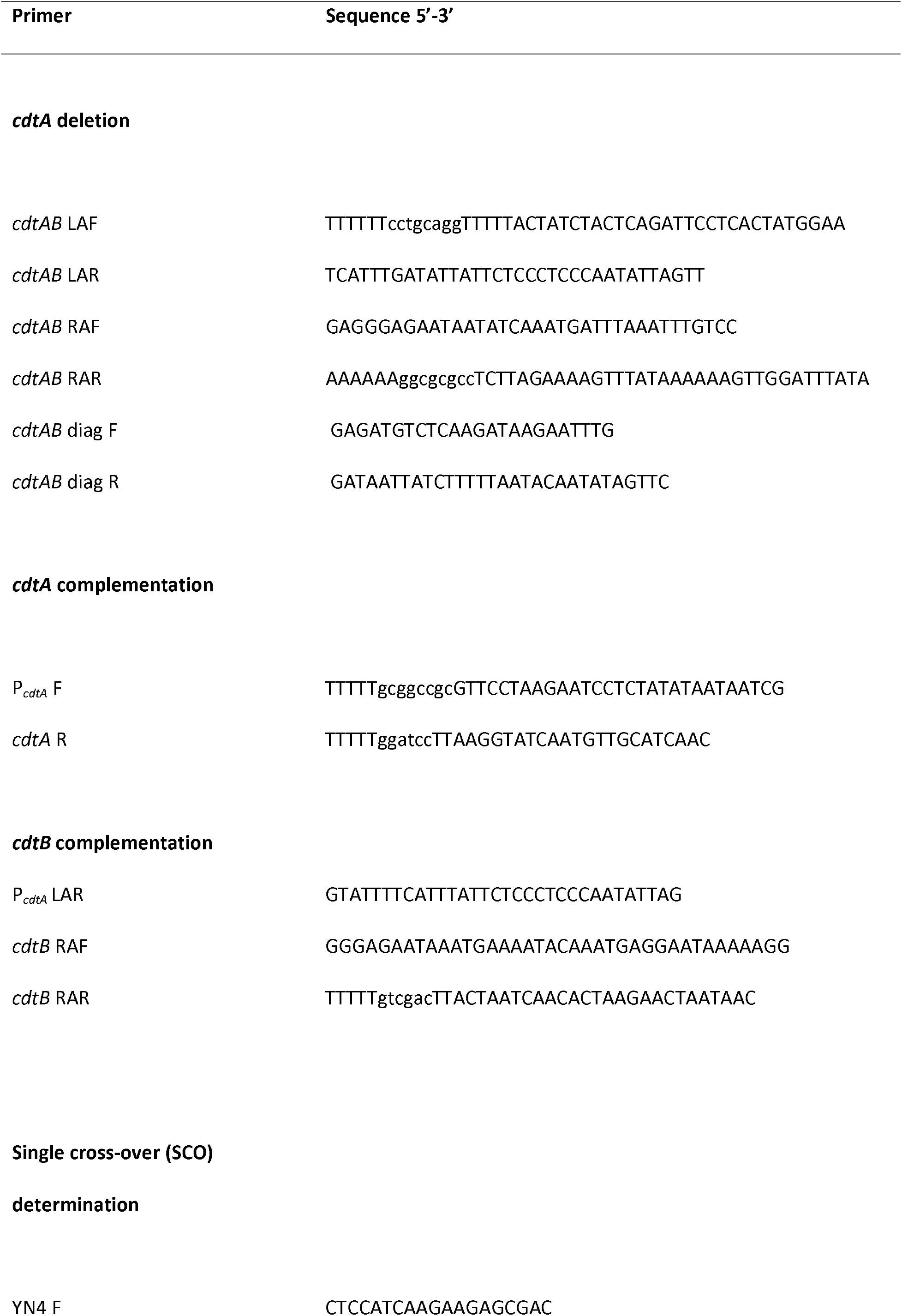
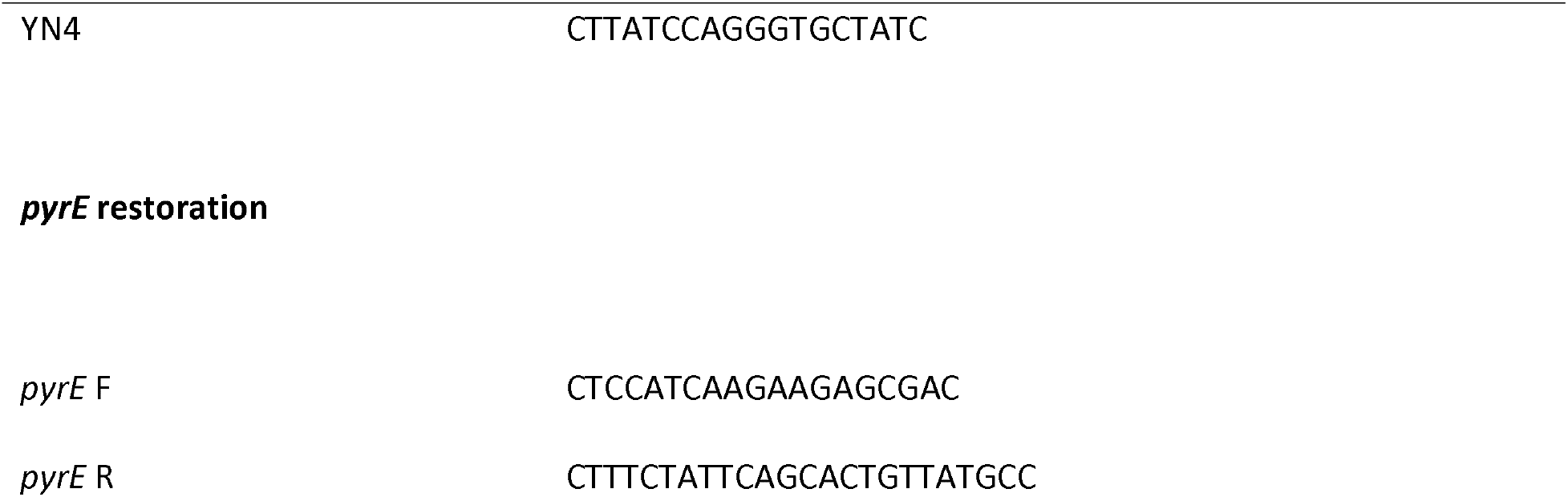
Oligonucleotide primers used in this study.

Strains differentially producing CDTa or CDTb, were generated by the integration of either *cdtA* or *cdtB* at the *pyrE* locus, under the control of *cdtA* promoter P_cdtA_. Firstly, *cdtA* coupled with its native promoter, was amplified by PCR using *P_cdtA_* F and *cdtA* R primers. The product of which was cloned into pMTL-YN2C by means of flanking *Not*I-*Bam*HI restriction sites thus generating the complementation cassette (CS) pMTL-YN2C-*P_cdtA_-cdtA.* In a similar fashion, pMTL-YN2C-*P_cdtA_-cdtB*, was generating by amplifying *P_cdtA_* using *P_cdtA_* F/P_*cdtA*_ LAR primers and *cdtB* using *cdtB RAF/cdtB* RAR primers, before SOEing the products together and cloning them into pMTL-YN2C by means of flanking *Not*I-*Sal*I restriction sites. Unfortunately, the CDTb-encoding construct could only be generated with an SNP in the promoter region of *P_cdtA_* ensuing an A-G substitution at position −124 relative to the start codon. The resultant plasmids were applied in parallel, to individually integrate the respective CDT constructs at the *pyrE* locus of *R20291*Δ*pyrE*Δ*cdtAB* concomitant with the repair of *pyrE*, following successful conjugation and selection for uracil prototrophs on CDMM lacking uracil. PCR analysis using primer *pyrE* WT F, coupled with either *cdtA* R or *cdtB* RAR, demonstrated effective knock-in at the *pyrE* locus (Figure 2B), thus generating strains *R20291*Δ*cdtAB*\*P_*cdtA*_-*cdtA* and *R20291*Δ*cdtAB*\*P_*cdrA*_-*cdtB*

### Analysis of CDT production by Western blot

Secreted CDTa/b was assessed by Western blot analysis of 48h culture-free supernatants exactly as described previously (13), using an HRP-Chicken *anti-Clostridium difficile* Binary Toxin Subunit A or B antibody (Gallus-Immunotech, USA).

### *C. difficile* Spore Preparation and Bacterial Culture

*C. difficile* spore stocks were generated as described previously (15). Briefly, *C. difficile* strains were grown in 2 mL of Columbia broth overnight at 37 °C anaerobically. The 2 mL inoculum was then added to 40 mL of Clospore media. The culture was incubated anaerobically at 37 °C for 5-7 days. Following the incubation, spores were harvested by centrifuging the culture at 3200 rpm for 20 min at 4 °C, then resuspending in cold sterile water. After washing the spores at least three times, the spore stocks were stored at 4 °C in sterile water. The stocks were heat treated at 65 °C for 20 min to eliminate any remaining vegetative cells. The concentration of spores in each stock was determined by serially diluting the stocks in anaerobic PBS and plating on BHI agar supplemented with 1% sodium taurocholate. Once the CFU/mL of each stock was determined, the infection inoculum was prepared by diluting the appropriate *C. difficile* strain spore stock to the appropriate concentration. Animals received 100 μL of inoculum each via oral gavage. To determine *C. difficile* colonization in infected animals, cecal contents were resuspended and serially diluted in reduced PBS. Serial dilutions were plated on BHI agar supplemented with 1% sodium taurocholate, 1 mg/mL cycloserine, and 0.032 mg/mL cefoxitin (Sigma), then incubated at 37 °C overnight in an anaerobic chamber. Bacterial burden was normalized to cecal content sample weight.

### Mice and *C. difficile* Infection

Experiments were carried out using 8 to 12-week-old male C57BL/6J mice from the Jackson Laboratory. All animals were housed under specific-pathogen free conditions at the University of Virginia’s animal facility, and all procedures were approved by the Institutional Animal Care and Use Committee at the University of Virginia. Mice were infected using a previously established murine model for CDI (10). Six days prior to infection, mice were given an antibiotic cocktail within drinking water consisting of 45 mg/L vancomycin (Mylan), 35 mg/L colistin (Sigma), 35 mg/L gentamicin (Sigma), and 215 mg/L metronidazole (Hospira). Three days later, mice were switched to regular drinking water for 2 days and the day prior to infection, given a single intraperitoneal injection of 0.016 mg/g clindamycin (Pfizer). The day of infection, mice were orally gavaged with vegetative (1 × 10 ^8^ CFUs) or spores (1 × 10^3^) of *C. difficile* strains as indicated. Mice were monitored daily during the course of infection and twice daily during the acute phase (days 2 and 3). Mice were immediately euthanized following the development of severe illness as measured by clinical scoring parameters. These parameters included weight loss, coat condition, eye condition, activity level, posture, and diarrhea, which were evaluated to give a clinical score between 0 and 20.

### Tissue Protein

Animals were humanely euthanized and cecal tissue was immediately removed. For cecal tissue lysate, cecal tissue samples were removed and washed with PBS. Lysing matrix (mpbio) and 400 μL of lysis buffer I (1X HALT protease inhibitor cocktail (Thermo Scientific), 5 mM HEPES (Gibco)) was added and tissue was bead beat for 1 min. 400 μL of lysis buffer II (1X HALT protease inhibitor cocktail (Thermo Scientific), 5 mM HEPES (Gibco), 2% Triton X-100 (Sigma)) was added and samples were gently inverted. Following a 30 min incubation on ice, samples were centrifuged at 13,000 xg for 5 min at 4 °C and supernatants were removed into a fresh tube. Samples were stored at −80 °C for later analysis.

### Hamsters and *C. difficile* Infection

Experiments were carried out using 90-100 g adult male Syrian Golden hamsters from Charles River Laboratory. All animals were housed under specific-pathogen free conditions at the University of Virginia’s animal facility, and all procedures were approved by the Institutional Animal Care and Use Committee at the University of Virginia. Hamsters were infected using a previously established hamster model for CDI (16), with minor modifications. Briefly, hamsters were orally gavaged with 0.03 mg/g clindamycin (Pfizer) five days prior to infection. On the day of infection, hamsters were orally gavaged with 10^2^ spores of *C. difficile* mutant strains as indicated. Hamsters were monitored twice daily over the course of infection, and were euthanized immediately upon development of severe illness as assessed via clinical scoring. Parameters used included weight loss, coat condition, eye condition, activity level, posture, and diarrhea, which were evaluated to give a clinical score between 0 and 20.

### Statistical Analysis

For animal work, survival curves were generated using the Kaplan-Meier estimator. Significance between groups was determined using ordinary one-way ANOVA, while Tukey’s test was used for multiple comparisons. Comparisons between two groups were done using a two-tailed *t* test. All statistical analyses were performed using GraphPad Prism software.

## Results

### CDTb induces TLR2/6-mediated NF-κB activation and IL-1β transcript expression

Our group has previously shown that CDT is recognized by TLR2/6 to induce downstream NF-κB activation (11). Considering that CDT consists of two components, CDTa and CDTb, we asked whether both were required to signal through TLR2/6, or if one component alone was sufficient. To test this, we utilized the HEKBlue-hTLR2 reporter cell line, which contains a transfected SEAP reporter for NF-κB, as well as human CD14/TLR2. Cells were treated with CDT holotoxin, CDTa, or CDTb and assessed for NF-κB activation as measured by secreted alkaline phosphatase (Figure 1A). Both CDT holotoxin and CDTb induced significant NF-κB activation, while CDTa did not. To investigate whether CDTb was activating NF-κB through the TLR2/6 heterodimer, cells were incubated with neutralizing antibodies against TLR2, TLR1, and TLR6. Blocking TLR2 or TLR6 significantly abrogated NF-κB activation following CDTb treatment, while blocking TLR1 had no effect. This indicates that CDTb is capable of inducing NF-κB activation in a TLR2/6-dependent manner.

**Figure 1:**
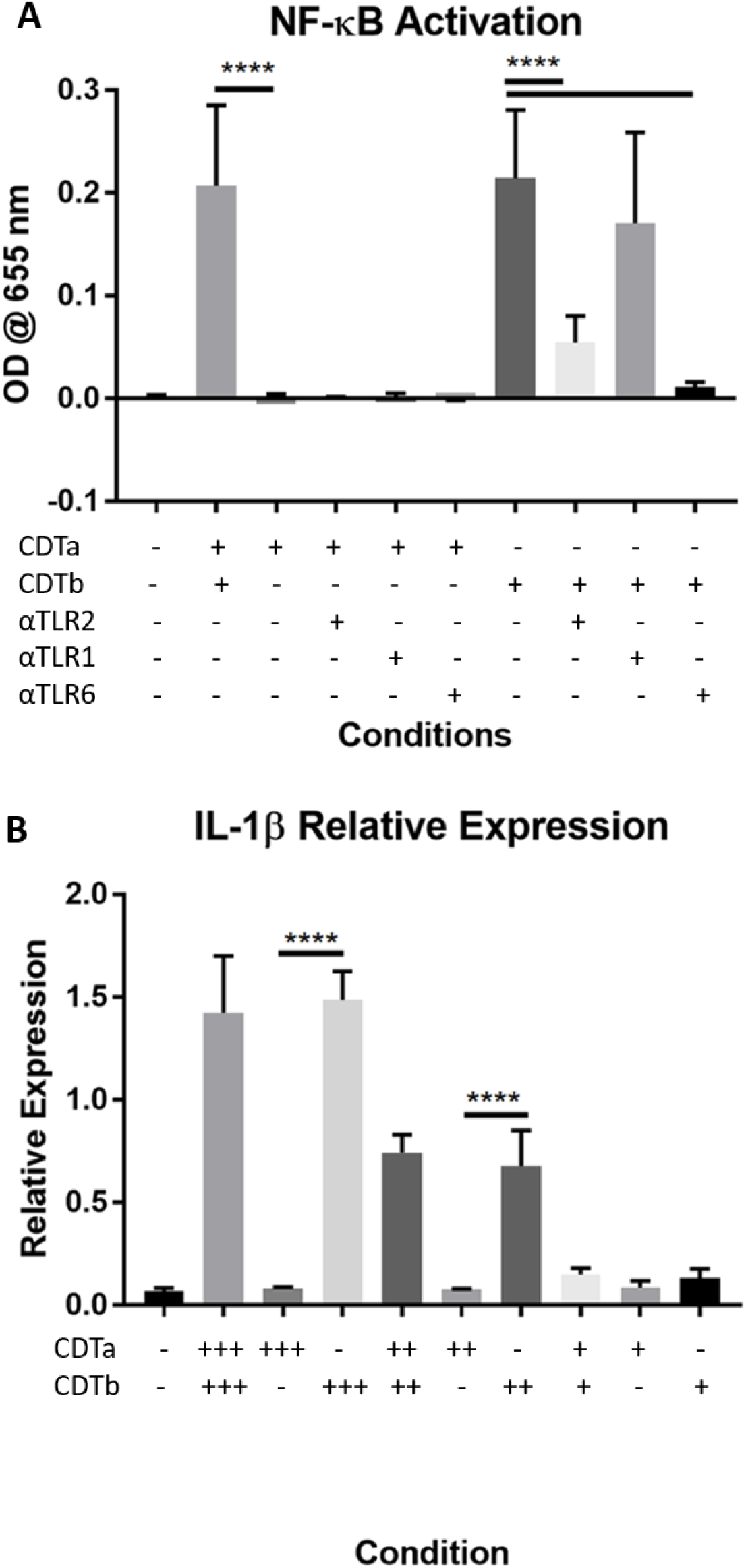
CDTb is capable of inducing NF-κB activation and IL-1β transcript expression. A) HEKBlue-hTLR2 cells were incubated at 37 C for 1 hour with 5 μg/mL of neutralizing antibodies against TLR2, TLR1, or TLR6, then treated with 5 ng/mL of CDTa and/or CDTb. Following an 8 hour incubation at 37 C, NF-κB activation was detected by measuring SEAP levels in the culture media via spectrophotometer. * * * * = p<0.0001 B) BMDCs were treated with decreasing amounts of CDTa and or CDTb as indicated for 18 h. +++200 ng/mL, ++20 ng/mL, +2 ng/mL. IL-1β transcript expression was assessed by RT-qPCR and was normalized to the Hprt housekeeping gene.****= p<0.0001

**Figure 2:**
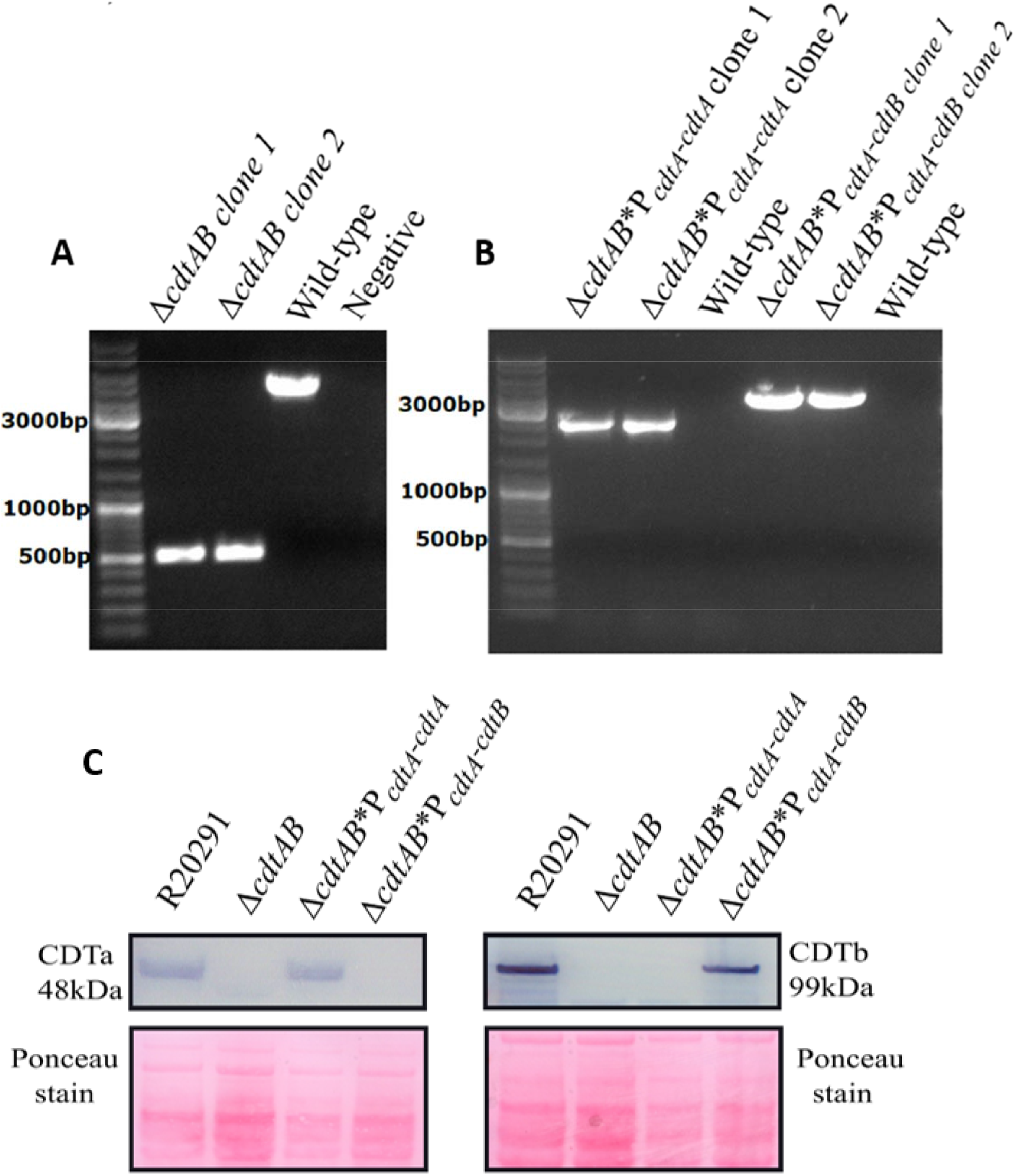
Authentication of *C. difficile* CDT mutant strains. Gel image following PCR analysis for A) deletion of *cdtAB* using *cdtAB* diag F/R primers and B) individual complementation of *cdtA* or cdtB at *pyrE* using *pyrE* WT F with *cdtA* R or *cdtB* R respectively. Gels ran alongside a GeneRuler DNA Ladder Mix (Thermo, USA). C) Western blot analysis of secreted CDT for 48 h culture-free supernatants detected with an HRP-Chicken *anti-Clostridium difficile* Binary Toxin Subunit A or B antibody. Ponceau staining was performed immediately following protein transfer and before blocking to ensure equal loading/transfer.

Next, we asked whether CDTb could induce inflammation in a more physiologically relevant cell system. Bone marrow-derived dendritic cells (BMDCs) from C57BL/6J mice were treated with CDT holotoxin, CDTa, or CDTb in decreasing concentrations and IL-1β transcript expression was measured via RT-qPCR (Figure 1B). Treatment with CDT holotoxin and CDTb induced robust expression of IL-1β transcripts, while there was no induction following treatment with CDTa. Additionally, lower doses of toxin induced lower levels of IL-1β expression, indicating that this response is dose-dependent. Altogether, these results indicated that the binding component of CDT is capable of inducing an inflammatory cytokine response in an immune cell type.

### Generation of CDT mutant strains

To determine the contribution of CDTb to disease pathology *in vivo*, CDTa(+)CDTb(−) and CDTa(−)CDTb(+) strains of R20291 were generated as outlined in the Methods section. These strains would express the primary clostridial toxins Toxin A and Toxin B, but would be mutated to express CDTa without CDTb, or CDTb without CDTa. Deletion of *cdtAB* and complementation with *cdtA* or *cdtB* was confirmed using PCR analysis (Figure 2A-B). Following successful strain development, we validated their phenotype regarding CDT production. To do this, we cultured R20291, R20291Δ*cdtAB*, R20291ΔcdtAB*P*cdtA*-*cdtA*, and R20291Δ*cdtAB*\*P*cdtA-cdtB* in TY broth and at the 48h time point, assessed each strain for CDTa/b production by Western blot analysis of culture-free supernatants using antibodies developed against CDTa or CDTb (Figure 2C). Analysis of the Western blots demonstrated that the *cdtAB* deletion mutant was devoid of detectable CDTa or CDTb production. As expected, individual complementation of *cdtA* restored CDTa production whilst complementation of *cdtB* restored CDTb production, thus generating strains differentially expressing CDTa or CDTb. The strains will hereafter be referred to as CDTa+ and CDTb+.

### The binding component CDTb does not increase virulence in a mouse model of *C. difficile* infection

Because CDTb is capable of inducing inflammation *in vitro*, we asked whether this response contributes to disease *in vivo.* To test this, we utilized a previously published (10) mouse model of *C. difficile* infection (CDI) (Figure 3A). We infected adult C57/BL6 mice with 1 × 10^3^ spores of either the R20291 wildtype strain, the Δ*cdtAB* strain (lacking CDTa and CDTb), or the CDTb+ strain (expresses CDTb but not CDTa). Two days post-infection, the mice were sacrificed and cecal tissue and contents were harvested for analysis. At this time point, mice infected with the wildtype strain began to experience some weight loss (Figure 3B) and significantly more severe disease as measured by clinical scoring (Figure 3C). However, mice infected with either the Δ*cdtAB* or the CDTb+ strain did not show any signs of weight loss or significant disease. Additionally, infection with the wildtype strain induced significantly higher levels of cecal IL-1β and IL-6 compared to either mutant strain (Figure 3D-E). There was no significant difference in Toxin A and Toxin 8 production *in vivo* (Figure 3F), indicating that the difference in virulence seen in the mutant strains was not due to a deficiency in production of either of these primary clostridial toxins. We examined whether the differences in disease severity between the wildtype strain and the CDT mutant strains were due to differences in bacterial colonization, but there was no significant difference in bacterial burden (Figure 3G). We also infected mice with vegetative cells of each strain (Supplementary Figure 1A-D) and found that while the wildtype strain induced significant mortality, weight loss, and clinical scores, both the Δ*cdtAB* and CDTb+ strains were avirulent. Overall, this indicates that expression of CDTb without CDTa is not sufficient to increase virulence in a mouse model of CDI.

**Figure 3:**
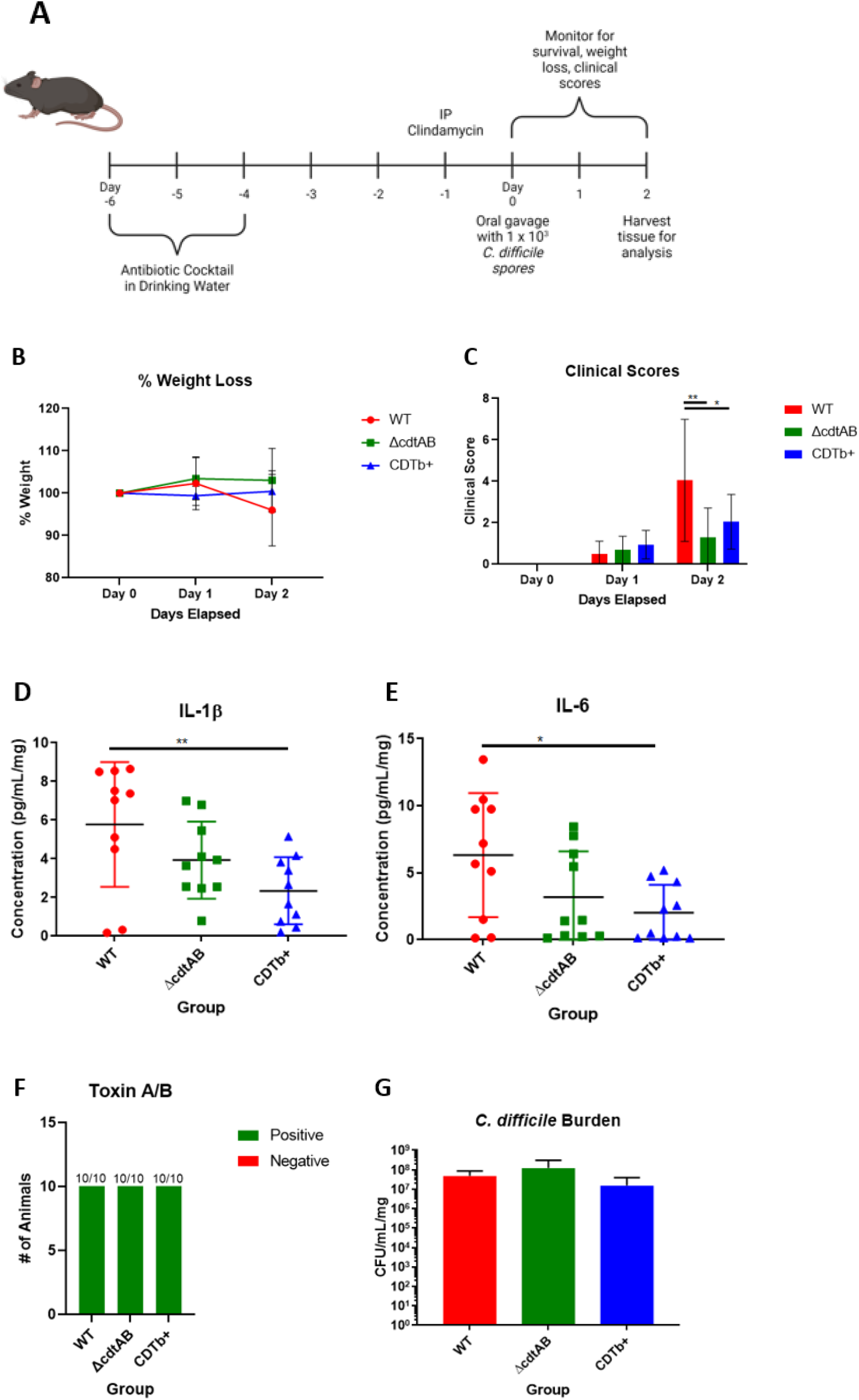
The binding component CDTb does not increase virulence in a mouse model of *C. difficile* infection. A) C57/BL6 mice were treated with antibiotics then infected with 1 × 10 ^3^ spores of R20291 wild-type (WT), R20291 Δ*cdtAB*, or R20291 CDTb+ strain of *C. difficile.* Following infection, mice were monitored for B) weight loss and C) clinical signs of disease (* * p=0.023, * p=0.0265). Data are combined from two independent experiments, n=20. Cecal cytokines D) IL-1β and E) IL-6 were measured by ELISA following cecal tissue lysis and are normalized to tissue weight. (D-E, **p=0.0098 and *p=0.0298) F) Presence of ToxinA/B in cecal contents was assessed via ELISA provided by TechLab. G) Cecal contents were suspended and serially diluted in anaerobic PBS, then plated on BHI agar with taurocholate and *C. difficile* supplement. Following overnight anaerobic incubation at 37 °C, colony growth was assessed and normalized to cecal content weight.

### The binding component CDTb enhances virulence in a hamster model of *C. difficile* infection

Despite the inability of the CDTb+ strain to induce disease in a mouse model, we hypothesized that the contribution of CDTb to disease pathology could be subtle, and as such may be overshadowed in the relatively resistant mouse model of CDI. Therefore, we asked whether a more sensitive animal model would reveal more minute differences in disease phenotype between infections with the mutant strains. To investigate this, we utilized a hamster model of CDI (Figure 4A). We infected adult Syrian Golden hamsters with 1 × 10^2^ spores of the Δ*cdtAB* strain or the CDTb+ strain, then monitored the animals twice daily for mortality (Figure 4B), weight loss, and clinical scores (Figure 4C). Infection with the CDTb+ strain induced significantly higher mortality as compared to the Δ*cdtAB* strain, and while the CDTb+-infected hamsters did experience more severe disease, the swiftness and severity of the infection prevented any statistically significant comparison between clinical scores. Indeed, the hamsters experienced a much more severe and much faster course of disease as compared to the mice, with the time between symptom onset and mortality being much shorter in the hamsters than in the mice. In a separate experiment, we sacrificed hamsters on either day one or day two post-infection and assessed levels of cecal IL-1β and IL-6 (Figure 4D-E), but saw no significant differences between hamsters infected with the Δ*cdtAB* strain and those infected with the CDTb+ strain, perhaps due to the rapidity of the disease course in this animal model. Overall, we find that the binding component of CDT increases virulence in a hamster model of CDI.

**Figure 4:**
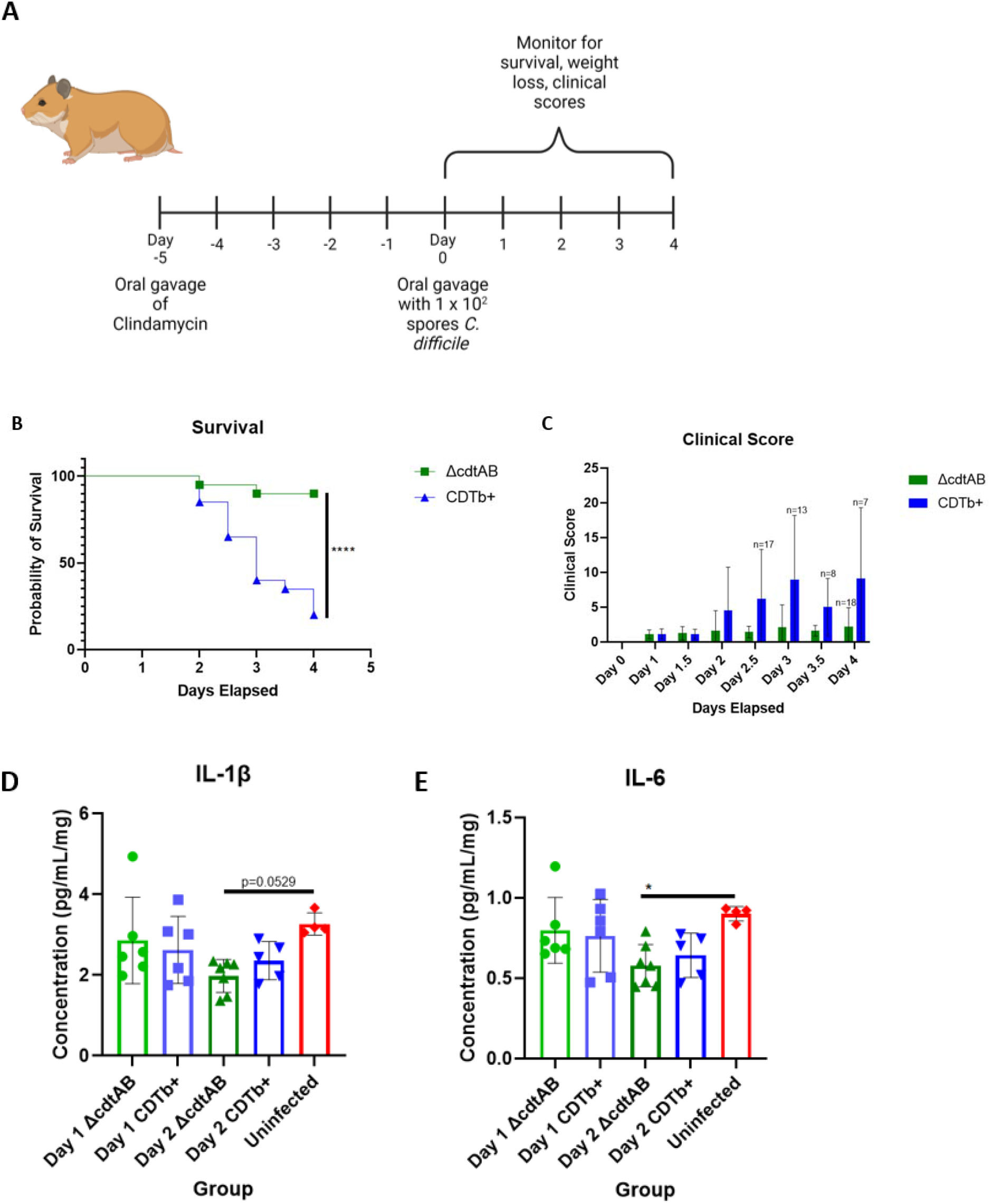
The binding component CDTb enhances virulence in a hamster model of *C. difficile* infection. A) Adult Golden Syrian hamsters were given oral clindamycin then infected with 1 × 10^2 spores of *C. difficile* R20291 Δ*cdtAB* or R20291 CDTb+. Following infection, hamsters were monitored for B) survival (****p<0.0001) and C) clinical signs of disease. Data were combined from two separate experiments, n=20. D, E) Hamsters infected with R20291 Δ*cdtAB* or R20291 CDTb+ were sacrificed on day 1 or day 2 post-infected and cecal cytokines were assessed via ELISA following cecal tissue lysis and are shown normalized to tissue weight. (*p=0.0389)

## Discussion

Our group has shown previously that the *C. difficile* binary toxin CDT is capable of activating the TLR2/6 heterodimer to induce downstream NF-κB activation (11). In this study, we further investigated this interaction and found that CDTb, but not CDTa, was sufficient to generate a TLR2/6-mediated NF-κB response. In addition, CDTb was capable of inducing a dose-dependent IL-1β response in BMDCs. This agrees with previous work from our group demonstrating that treatment of immune cells with CDT holotoxin stimulates an inflammatory response (10). The results shown here indicate that CDTb is capable of generating inflammation independently, outside of its established role in facilitating the entry of the enzymatic component, CDTa, into the host cytosol. Further supporting the idea that CDTb can act on cells independently of CDTa, it has recently been shown that CDTb can form pores in the cytoplasmic membrane and induce cell death in the absence of CDTa (17, 18). In addition, another member of the binary toxin family, iota toxin from *Clostridium perfringens*, has been shown to act similarly, as the binding component was capable of inducing cell necrosis (19). Altogether, these studies demonstrate the capability of binary toxin binding components to act in the absence of their enzymatic components and highlight the need for further investigation into the scope of their contributions during infection.

While not common, clinical strains of *C. difficile* that are negative for the primary clostridial toxins TcdA and TcdB but positive for CDT have been isolated from symptomatic patients (20), supporting the idea that CDT does play a significant role during CDI. However, the individual contributions of CDTa and CDTb have been understudied. To investigate the role of CDTb during infection, we developed novel mutant strains of *C. difficile* that differentially express CDTa or CDTb at levels comparable to the wildtype parental strain. This allowed us to study the effect of CDTb expression on the development of disease. Thus, these strains represent valuable tools that can be used to study the effects of each CDT component in the full context of bacterial infection.

In this study, we saw that the CDTb+ mutant strain did not cause disease in a mouse model, but did induce severe disease and mortality in a hamster model of infection. This striking difference suggests that there may be entirely different mechanisms responsible for driving disease severity and mortality between these models. Further supporting this idea are the marked differences in disease progression we observed. In the mouse model, the animals experienced a more prolonged course of disease and were capable of potentially recovering back to their baseline weight and appearance. In the hamster model, however, the animals underwent a very rapid and severe course of infection, in some cases progressing from no outward signs of disease to moribund in less than six hours. In addition, no recovery took place for any animal once they developed symptomatic disease. It is unclear what is responsible for such a high degree of variation in disease progression. One possible difference may be in the kinetics of bacterial colonization, as more rapid or more gradual colonization could be responsible for the contrasting disease progression. The extent of epithelial barrier disruption may also play a role, as differences in toxin receptor expression could affect the degree of damage induced by the toxins. Similarly, innate immune receptor expression may influence the scale of the inflammatory response induced in response to infection. These all represent potential avenues for further investigation into the differing responses between these animal models.

While we were able to determine that the CDTb+ strain was more virulent in the hamster model, it is not yet known how exactly CDTb is contributing to worsened disease. It may be that hamsters express higher levels of TLR2 and/or TLR6, and as such experience greater levels of TLR2/6-mediated inflammation. Another possibility is that hamsters are more sensitive to epithelial damage induced by the cytotoxic activity of CDTb (17, 18). Further study into these mechanisms can help shed light on the effects of CDTb during infection.

Overall, we have found that the binding component of the *C. difficile* binary toxin is capable of inducing inflammation and contributes significantly to disease in a hamster model. To our knowledge, this is the first work demonstrating the impact of CDTb toxicity *in vivo* and helps to further explain the heightened virulence displayed by the epidemic strains of *C. difficile.* Understanding the significance and impact of CDT and its individual components during infection can help in developing therapeutic strategies against these more severe hypervirulent strains.

## Declaration of competing interest

WAP Jr. is a consultant for TechLab, Inc.

## Acknowledgements

*C. difficile* Toxin A/B ELISA kits were provided by TechLab. This work was supported by the National Institutes of Health (grant numbers R0l A|124214 to WAP, T32AI007496 to MS, and F32DK124048 to JLL) and grants from the Marie Curie Clospore ITN, contract number 642068, and the NIHR Nottingham BRC (Reference no: BRC-1215-20003). The views expressed are those of the authors and not necessarily those of the NHS, the NIHR or the Department of Health.

## Author Contributions

MS designed, performed, and analyzed the data from the cell culture and animal experiments. JL, AD, JU, and WP helped with tissue processing and provided invaluable advice. NM constructed the initial *cdtAB* deletion mutant. TB repaired the *pyrE* allele, constructed the *cdtA/cdtB* complements and analyzed CDT production. SK and NPM supervised NM and TB. CS provided the purified CDT toxin. WAP Jr. supervised MS and supported all aspects of the work.

---

**Supplementary Figure 1:**
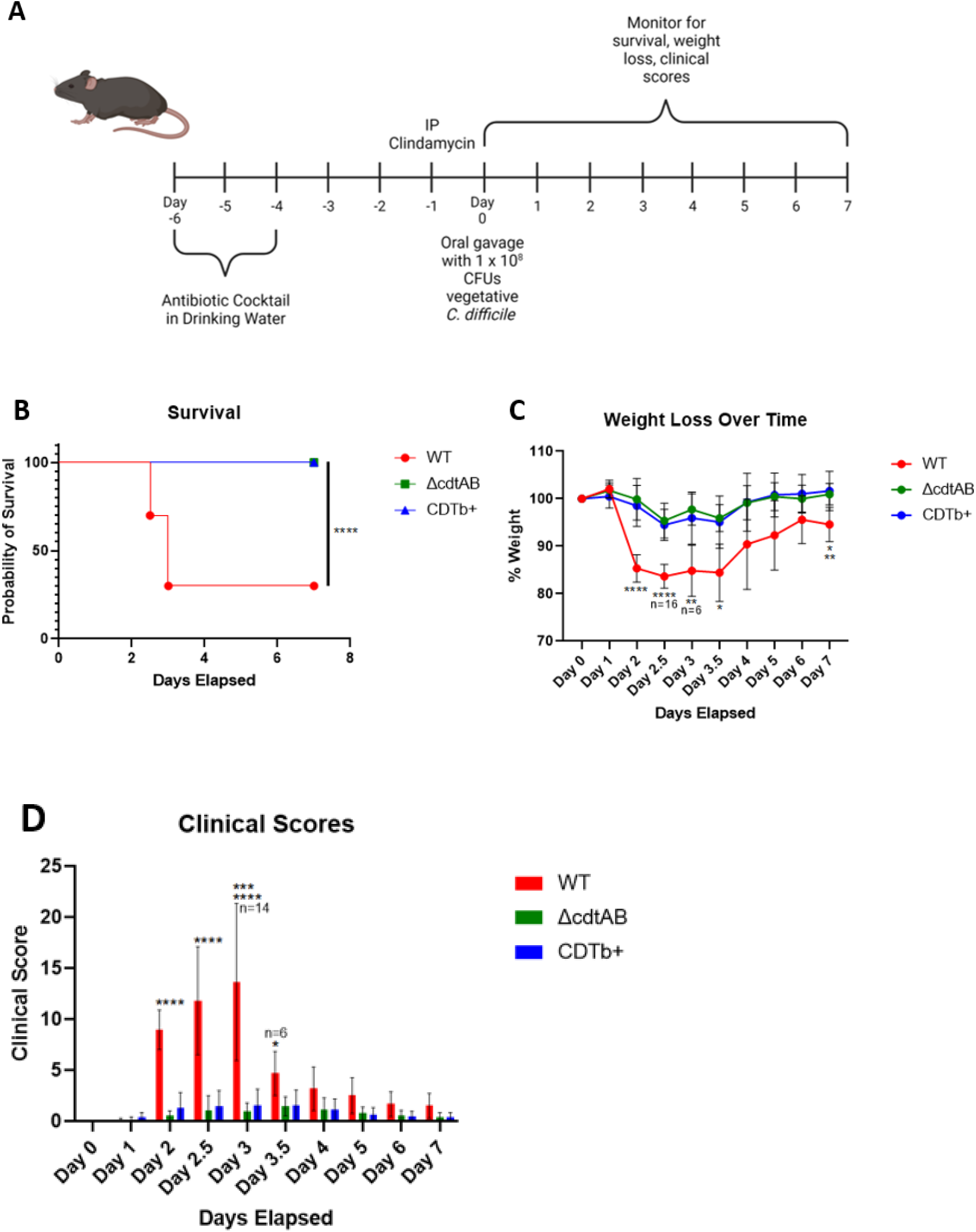
Vegetative infection with CDTb+ *C. difficile* strain. A) C57/BL6 mice were treated with antibiotics then infected with 1 × 10^8 CFUs of *C. difficile* R20291 wildtype (WT), R20291 Δ*cdtAB*, or R20291 CDTb+ strain. Following infection, mice were monitored for B) survival (* * * *p<0.0001), C) weight loss, (****p<0.0001, **p=0.032 (Δ*cdtAB)*, * * p=0.051 (CDTb+), *p=0.0111 (Δ*cdtAB)*, *p=0.0129 (CDTb+), * p=0.0138 (Δ*cdtAB)*, ** p=0.0066 (CDTb+)) and D) clinical signs of disease (****p<0.0001, ** *p=0.0001, *p=0.0312 (Δ*cdtAB)*, *p=0.0338 (CDTb+). Data were combined from two independent experiments, n=20.

## Notes

### Competing Interest Statement

William A. Petri Jr. is a consultant for TechLab, Inc.

